# Drug Target Identification and Virtual Screening in Pursuit of Phytochemical Intervention of *Mycobacterium chelonae*

**DOI:** 10.1101/315408

**Authors:** Zarrin Basharat, Shumaila Zaib, Azra Yasmin, Yigang Tong

## Abstract

*Mycobacterium chelonae* is a rapidly growing mycobacterium present in the environment. It is associated with skin and soft tissue infections including abscess, cellulitis and osteomyelitis. Other infections by this bacterium are post-operative/transplant-associated, catheter, prostheses and even concomitant to haemodialytic procedures. In this study, we employ a subtractive genomics approach to predict the potential therapeutic candidates, intended for experimental research against this bacterium. A computational workflow was devised and executed to procure core proteome targets essential to the pathogen but with no similarity to the human host. Initially, essential *Mycobacterium chelonae* proteins were predicted through homology searching of core proteome content from 19 different bacteria. Druggable proteins were then identified and N-acetylglucosamine-1-phosphate uridyltransferase (GlmU) was chosen as a case study from identified therapeutic targets, based on its important bifunctional role. Structure modeling was followed by virtual screening of phytochemicals (N > 10,000) against it. 4,4’-[(1E)-5-hydroxy-4-(methoxymethyl)-1-pentene-1,5-diyl]diphenol, apigenin-7-O-beta-gluconopyranoside and methyl rosmarinate were screened as compounds having best potential for binding GlmU. Phytotherapy helps curb the menace of antibiotic resistance so treatment of *Mycobacterium chelonae* infection through this method is recommended.

## INTRODUCTION

Mycobacteria are categorized into two major groups, tubercular and non-tubercular mycobacteria. Nontuberculous mycobacteria are further divided into two groups, rapidly growing and slow growing mycobacteria, depending upon duration of their reproduction in suitable medium (Tortoli, 2014). *Mycobacterium chelonae* is a member of rapidly growing group, which takes less than a week for its reproduction in/on medium. It is the most commonly isolated organism among all rapidly growing mycobacteria. It is mostly found in water sources and on medical instruments such as bronchoscopes (Gonzalez-Santiago and Drage, 2015), and has been isolated from environmental, animal and human sources. *Mycobacterium chelonae* infections in human hosts have increased over time, with reports of both haematogenous and localized occurrences in the recent past (Hay, 2009).

Outbreaks due to *Mycobacterium chelonae* contaminated water and compromised injections are a rising problem. Infections have been linked to cosmetic and surgical procedures, such as trauma, surgery, injection (botulinum toxin, biologics, dermal fillers), liposuction, breast augmentation, under skin flaps, laser resurfacing, skin biopsy, tattoos, acupuncture, body piercing, pedicures, mesotherapy and contaminated foot bath (Gonzalez-Santiago and Drage, 2015). *Mycobacterium chelonae* may also colonize skin wounds, as result of which patients form abscess, skin nodules and sinus tracts (Patnaik *et al.*, 2013).

Recent era has observed a trend for search of drug targets in pathogens using computational methods, with a focus on genomic and proteomic data (Shanmugham and Pan, 2013).

Comparative/differential and subtractive genomics along with proteomics has been used by many researchers for the identification of drug targets in various pathogenic bacteria like *Pseudomonas aeruginosa, Helicobacter pylori*, *Brugia malayi*, *Leptospira interrogans*, *Listeria monocytogenes*, *Mycobacterium leprae* (Amineni *et al.*, 2010; Shanmugam and Natarajan, 2010; Sarangi *et al.*, 2015) etc. Determination of potential drug targets has been made possible by the availability of whole genome and their inferred protein complement sequences in public domain databases (Sarangi *et al.*, 2015). Numerous anti-tubercular drug targets, even though mostly for *Mycobacterium tuberculosis* have been reported previously. Some of these include proteins essential for survival or the ones that could be targeted by anti-microbial peptides. Common ones include sulphur metabolic pathway proteins (Reviewed by Bhave *et al.* 2007) and folate pathway proteins (Rengarajan *et al.*, 2004) but due to mutations, folate pathway proteins are prone to drug resistance as well.

Parenteral antibiotics against *Mycobacterium chelonae* include tobramycin, amikacin, imipenem, and tigecycline, but it has demonstrated resistance to antibiotics and disinfectants. This property assists *Mycobacterium chelonae* in colonizing water systems and allows its access to humans (Jaén-Luchoro *et al.*, 2016). Successful resolution of *Mycobacterium chelonae* infection with antibiotics has been reported in different scenarios (Brown-Elliott *et al.*, 2001; Choi *et al.*, 2018), but resistance to beta-lactams (Jarlier *et al.*, 1991) and multiple drugs (Churgin *et al.*, 2018) has also been observed. Till now, no specific guidelines for the treatment of *Mycobacterium chelonae* have been defined in the literature (Gonzalez-Santiago and Drage, 2015). Antibiotic resistance also illustrates the need to search out new drug targets for design of better therapies against infection by this bacterium, especially using naturally existing metabolites from microbes and plants. In the current study, subtractive proteomics was applied to identify essential druggable proteins in *Mycobacterium chelonae*. Docking of the selected protein GlmU with phytochemicals was carried out for identification of a candidate which might bind it and stop its normal cellular function, leading to bacterial lysis/death.

## MATERIAL AND METHODS

### Prediction of *Mycobacterium chelonae* essential proteome

Complete proteome sequence of *Mycobacterium chelonae* CCUG 47445 with accession no: **<u>NZ_CP007220</u>** was downloaded from the NCBI database. For the prediction of essential proteins, Geptop (Wei *et al.*, 2013) was installed on computer and a search was carried out to align the *Mycobacterium chelonae* protein sequences against the essential or core protein sequences from defined set of 19 bacteria with an essentiality score cut-off value range of 0.15 (Wei *et al.*, 2013). Results were saved and analyzed further according to the workflow (Fig. 1).

**Fig. 1.**
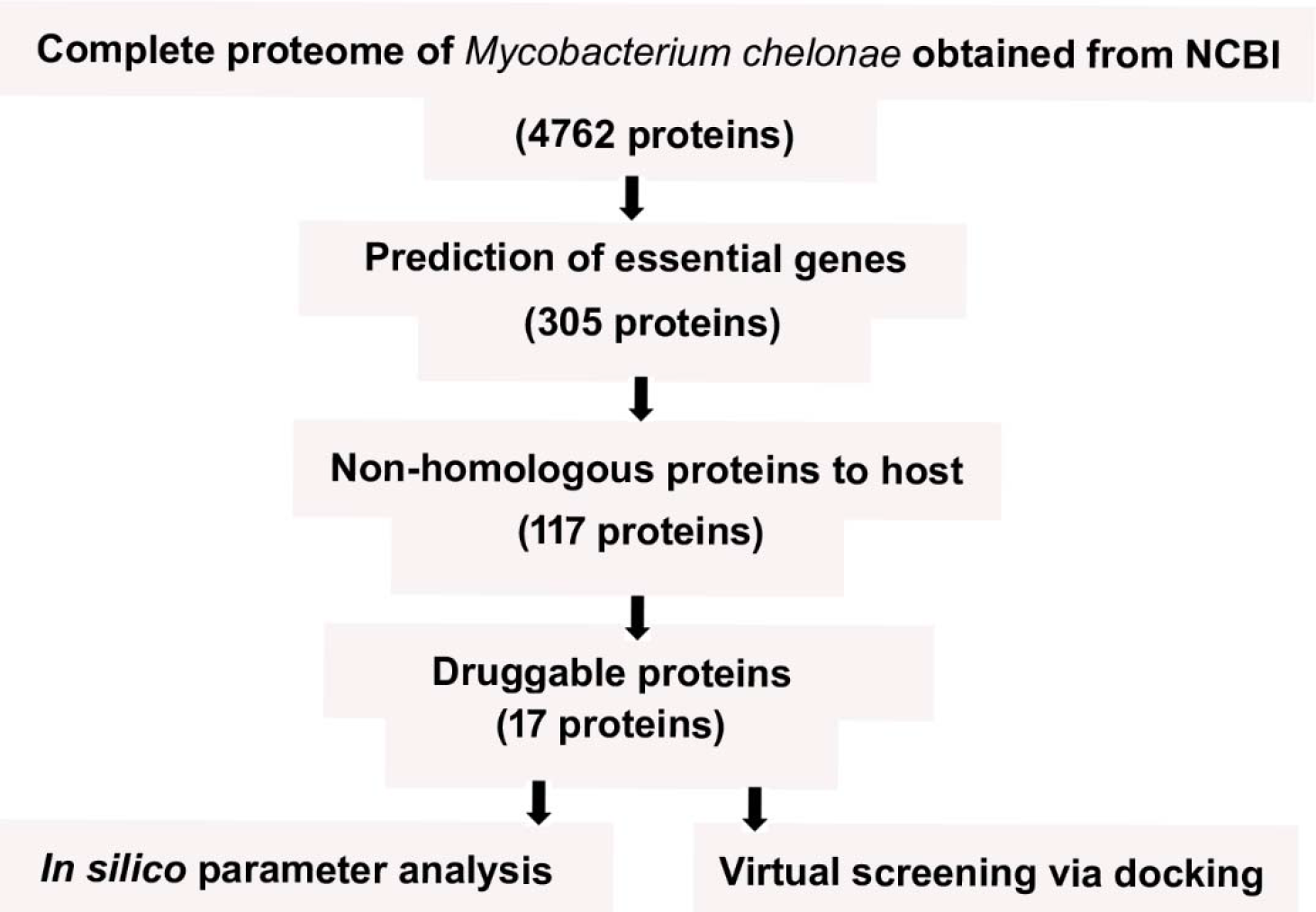
Workflow describing subtractive proteomics with respective number of obtained sequences. *In silico* parameter analysis included hydrophobicity, sub-cellular location prediction, molecular weight etc.

### Prediction of non-homologous host proteins

In order to find the bacterial proteins which do not have similarity with human host, the set of essential protein sequences of *Mycobacterium chelonae* was subjected to BLASTP against the human proteome database (Uniprot release 2014). The standalone BLAST software was used for this purpose. For identification of non-homologous proteins, an expectation (E-value) cut-off of 10^−2^, gap penalty of 11 and gap extension penalty of 1 was set as the standard. E-value cut-off (10^−2^), based on reported research protocols (Sarangi *et al.*, 2015) was considered.

### Identification of putative drug targets

There are several molecular and structural properties which have been explored by researchers for selecting suitable therapeutic targets in pathogenic microorganisms. These properties include determination of molecular weight, sub-cellular localization, 3D structure and druggability analysis. These properties were evaluated for selection of the therapeutic targets in *Mycobacterium chelonae*. Molecular weight was calculated by using computational tools and drug target-associated literature available in the Swiss-Prot database. Subcellular localization of therapeutic targets was predicted using PSORTb (Nancy *et al.*, 2010). It uses feature support vector machine-based method and suffix tree algorithm for downstream analysis. Predictions are grouped through a Bayesian scheme into one final (consensus) result. Druggable targets were identified with BLAST hits through unified protocol from the DrugBank. Parameters were: gap cost: −1 in case of extension or opening, mismatch penalty: −3, E-value: 1* 10^−5^, match: 1, filter algorithm: DUST and SEG (Azam and Shamim, 2014). The targets were subjected to KEGG blast for identification of associated pathways.

### Virtual screening of ligand against selected target

Keeping in view the results of sub-cellular localization, molecular weight determination and druggability analysis, an essential protein (GlmU) was chosen for further downstream processing. Swiss Model was used for the prediction of 3D structure of the selected target protein (Biasini et al., 2014). This tool constructs structure model by recognizing structural templates from the PDB using multiple threading alignment approaches (Wang *et al.*, 2016). The top structure used for structure prediction was that of GlmU from *Mycobacterium tuberculosis* (PDB ID: 3D8V). The structure was validated and analyzed for quality using SAVES (https://services.mbi.ucla.edu/SAVES/), consisting of ERRAT, VERIFY3D and Ramachandran plot analysis. TM-score was calculated using TM-align (Zhang and Skolnick, 2005).

Phytochemical libraries consisting of three groups were downloaded and docked with GlmU. group I consisted of ~2000 ayurvedic pharmacopia compounds (downloaded from http://ayurveda.pharmaexpert.ru/) (Lagunin et al., 2015), group II had ~6000 North African flora compounds (Ntie-Kang et al., 2017) (downloaded from http://african-compounds.org/nanpdb/) and group III had 2266 phytochemical compounds from various other medicinal plants (available from http://bioinform.info/in.sdfformat) (Mumtaz et al., 2016). Blind docking was carried out using Molecular Operating Environment (MOE) with the parameters: placement: triangle matcher, rescoring 1: London dG, refinement: forcefield, rescoring 2: affinity dG. MOE provides fast and accurate docking results based on dedicated algorithms and accurate scoring functions. Structural preparation program embedded in MOE added the missing hydrogen atoms, corrected the charges and assigned near precise hybridization state to each residue (Basharat *et al.*, 2017). Compounds with least energy and fulfilling the criteria of Lipinski’s drug like test were considered for further analysis. ADME/Tox analysis was also performed for the top compounds by inputting the structure at the webserver https://preadmet.bmdrc.kr/.

## RESULTS AND DISCUSSION

### Essential proteome prediction

Initially, total proteome of *Mycobacterium chelonae* was subjected to core or essential proteins prediction. Geptop identified these proteins by screening against 19 bacteria (Supplementary Table 1), based upon orthology and phylogeny features. In the process of essential proteome mining through homology-based methods, a query protein is considered as essential if it is also present in another bacterium and experimentally identified as essential for survival. There are various methods for the prediction of essential proteins, for example single-gene knockout, transposon mutagenesis and RNA interference but all these methods are time consuming and laborious. A good alternative is high-efficiency computational methods designed specifically for this type of work (Wei *et al.*, 2013). Predicted essential proteins were 305 in number (Supplementary Table 2), linked with significant metabolic pathways in the pathogen life cycle and necessary for its survival. In order to disrupt the function and existence of pathogen it is most important to attack those bacterial proteins which regulate important functions (e.g. nutrient uptake) in the host environment (Butt *et al.*, 2012). Latest antimicrobial drugs are designed on the principle of the inhibition of the pathogen’s metabolic pathway. Therefore, such protein sequences may be considered as possible therapeutic targets.

### Identification of non-host proteins

Non-host proteins refer to those bacterial proteins which do not have homology with human proteins. If the homologous proteins are targeted, they could badly affect the metabolism of host due to similarity with host proteins. Therefore, non-host proteins could be preferred better drug targets, as side effects and cross-reactivity caused by the use of antibiotics could be evaded for harming host (Sarangi *et al.*, 2015). Among the core proteome of *Mycobacterium chelonae*, 117 proteins (Supplementary Table 3) indicated ‘no hit’ against the human proteome according to the set criteria. These proteins were then used for subsequent analysis.

### Drug Target mining and analysis

BLASTp was performed to identify significant drug target from newly selected essential proteins. Only 17 proteins had significant hits against druggable proteins present in the DrugBank (Table 1).

**Table 1.**
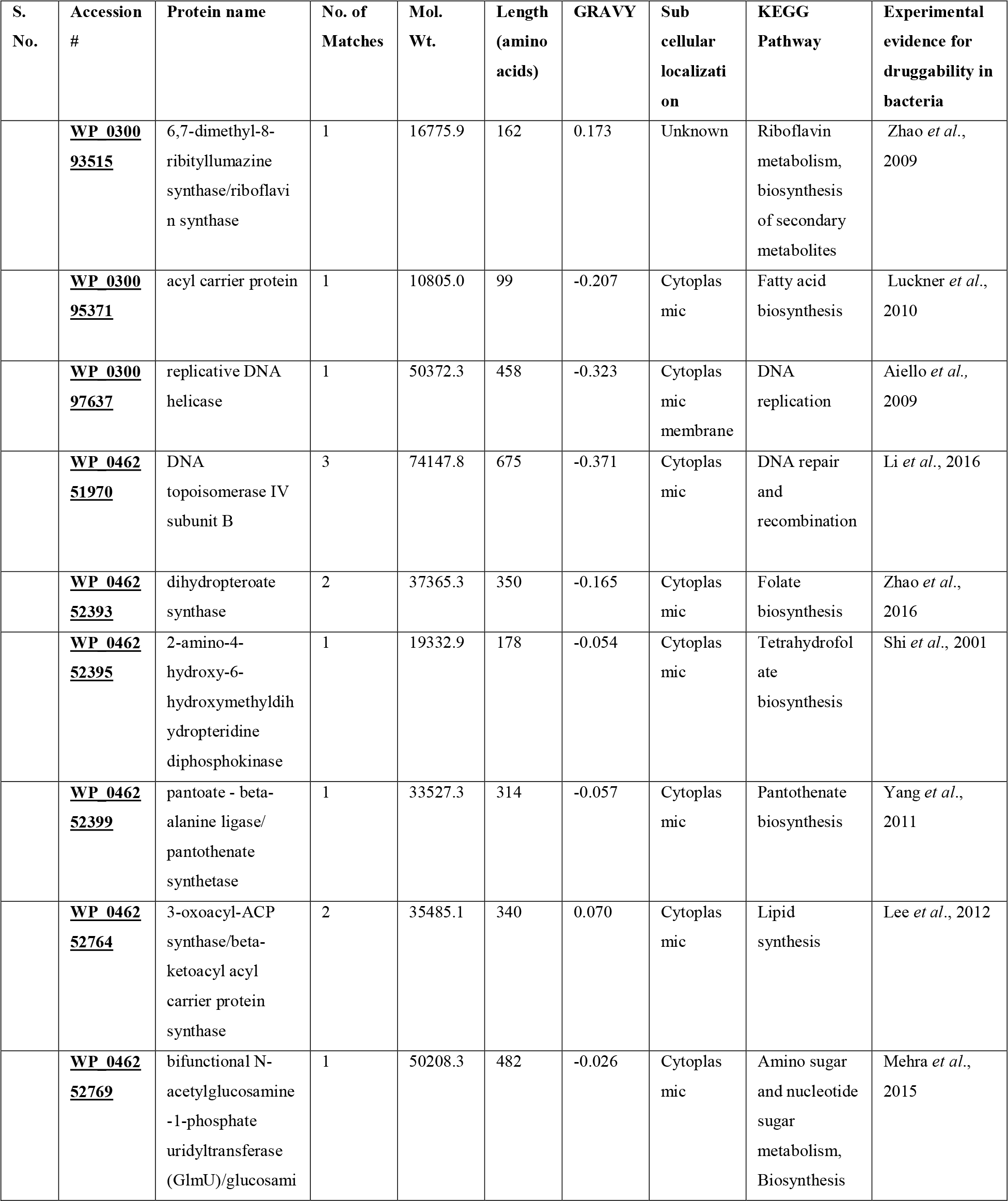
Predicted druggable proteome of *Mycobacterium chelonae.* GRAVY is average hydrophobicity of the protein measured by Kyte-Doolittle algorithm. Hydrophobicity value below 0 are indicates globularity (means protein is hydrophilic), while score value of above 0 indicates protein to be membranous (hydrophobic)

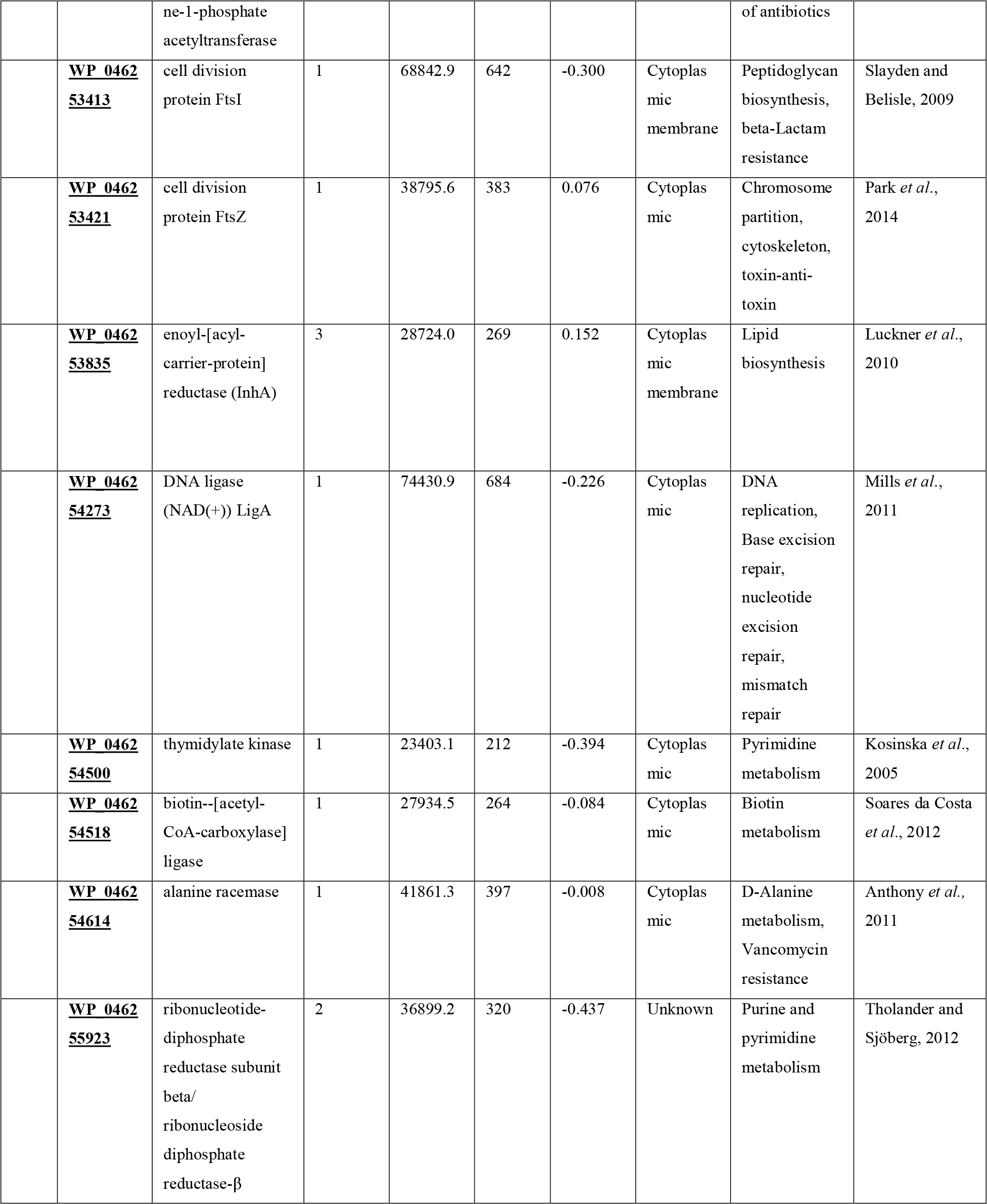

Molecular weight determination and druggability analysis could improve the screening process for therapeutic targets, as observed previously for numerous pathogenic bacteria and fungi (Abadio *et al.*, 2011). The molecular weight for each potential drug target was calculated (Table 1) and based upon previous studies, it is suggested that smaller proteins are readily soluble and easier to purify (Duffield *et al.*, 2010).

Sub-cellular localization is a critical factor as it helps in accessing the target gene. Cellular functions are compartment specific, so if the location of unknown protein is predicted then its function could also be known which help in selection of proteins for further study. Membrane proteins are reported as more useful target and more than 60% of the currently known drug targets are membrane proteins (Tsirigos *et al.*, 2015). Cytoplasm is the site of proteins synthesis and most of these proteins remain there to carry out their specific functions after synthesis. However, some proteins need to be transported to different cellular compartments for their specific function (Strzyz, 2016). The subcellular localization of non-host proteins of *Mycobacterium chelonae* was predicted and majority were demarcated as cytoplasmic (Table 1).

### GlmU analysis and phytochemical screening

After the characterization of all druggable proteins of *Mycobacterium chelonae*, GlmU was selected for further analysis (out of 305 essential and 117 non-homologous proteins). It is a bifunctional enzyme, exhibiting both acetyltransferase and uridyltransferase activities (Sharma *et al.*, 2016) and involved in biosynthesis of peptidoglycan. Peptidoglycan in complex with mycolyl-arabinogalactan is important for the viability of Mycobacteria as it upkeeps the basal structure necessary for upper “myco-membrane” support. GlmU has been analyzed for various bacterial species, including *Escherichia coli*, *Streptococcus pneumonia*, *Haemophilus influenzae* and *Mycobacterium tuberculosis* (Patin *et al.*, 2015). Predicted structure of GlmU had an estimated RMSD of 0.5 Å from the reference structure. A TM-score of 0.97 was obtained and since a score of 0.5-1 means same fold, with greater value meaning more similarity. 0.97 meant a very similar fold structure. Structure was found to be a homotrimer. Ramachandran plot showed 97.2% residues in favored regions (90.4% in core, 9.4% in allowed, 0.3% in generously allowed) and no residue in disallowed region (Fig. 2). According to VERIFY3D program, at least 80% of the amino acids should have value >= 0.2 in the 3D/1D profile and 96.55% of predicted GlmU residues had an averaged 3D-1D score >= 0.2, thus passing the quality check. An ERRAT score of 90.66 was obtained.

**Fig. 2.**
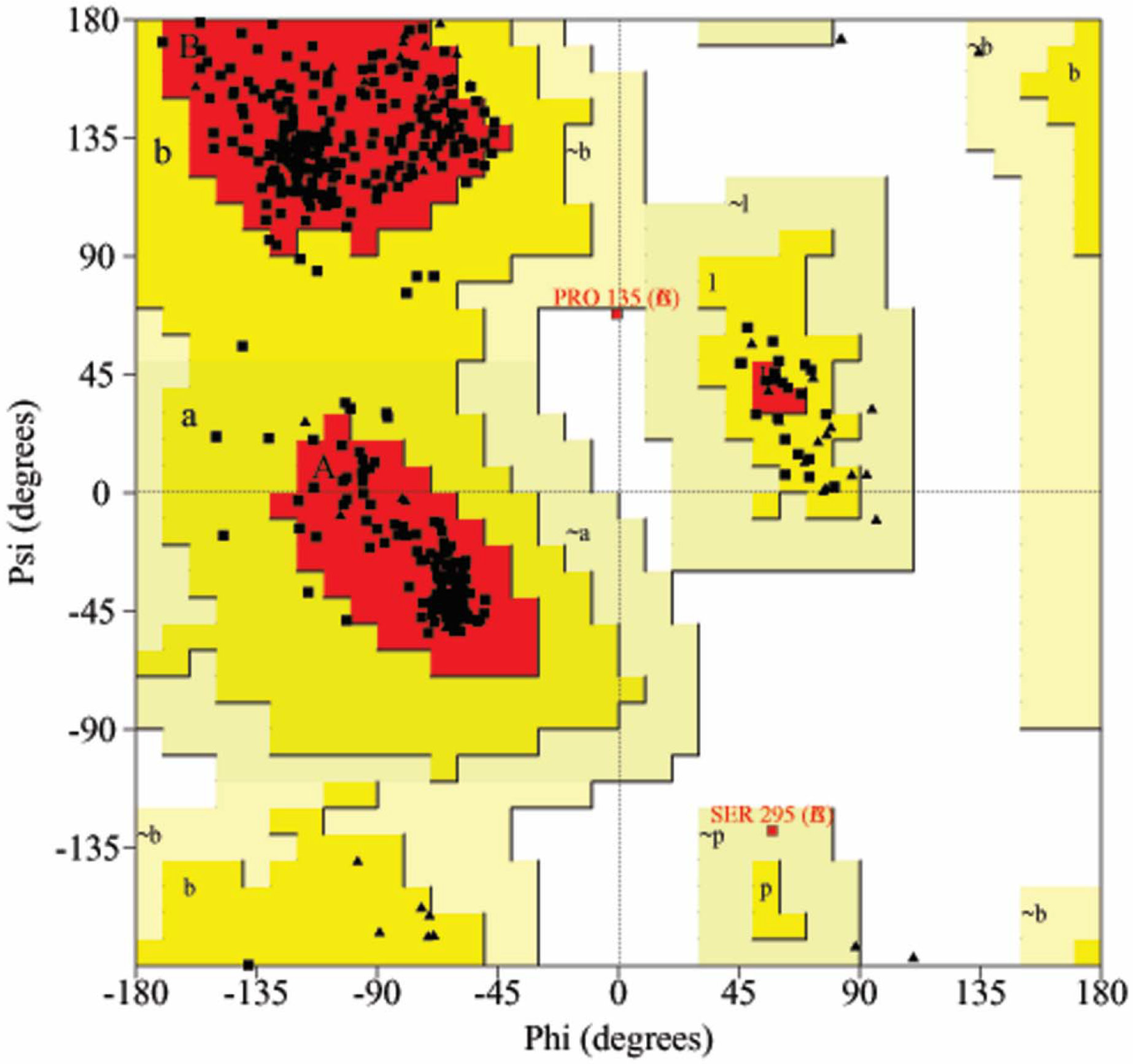
Ramachandran plot of the predicted GlmU structure.

Each monomer of GlmU consists of two domains: N and C-terminal domains. N-terminal domain has α/β like fold, similar to dinucleotide-binding Rossmann fold topology. C-terminal domain exhibits a regular left-handed β-helix conformation and a long α-helical arm connecting both domains (Sharma *et al.*, 2016). N-terminal domain is essential for uridyltransferase activity as it catalyses the transfer of uridine monophosphate from uridine-triphosphate to N-acetylglucosamine-1-phosphate (GlcNAc1P). C-terminal domain has acetyltransferase activity as it catalyzes the transfer of an acetyl group from acetyl-CoA coenzyme to GlcN1P, in order to produce GlcNAc1P (Moraes *et al.*, 2015).

GlmU plays fundamental role in the formation of bacterial cell wall by carrying out catalysis of uridine-diphospho-N-acetylglucosamine, an important precursor in bacterial peptidoglycan cell wall (Sharma *et al.*, 2016). Activity of acetyltransferase of eukaryotic cells differ from the activity of GlmU in the way that, eukaryotic acetyltransferase occurs on GlcN6P and not on GlcN1P. These properties make GlmU suitable as the drug target (Moraes *et al.*, 2015) and it has been used for drug targeting in bacteria such as *Haemophilus influenza* (Mochalkin *et al.*, 2007) apart from designing inhibitors against it for Mycobacterial species (Mehra *et al.*, 2016). It is also known that proteins that are involved in more than one pathway of pathogen, in addition to that they are non-host proteins, could be more effective drug targets (Sarangi *et al.*, 2015). Inactivation of bifunctional GlmU enzyme leads to loss of mycobacterial viability, therefore GlmU was used for docking against phytochemicals.

The possible interaction between protein and the ligand is understood computationally through molecular docking therefore receptor centric docking approach was employed for screening of prospective phytochemical library of compounds from more than 10,000 plant derived compounds against GlmU of *Mycobacterium chelonae*. Whole surface of the protein was scanned for best docking pose. A comparative analysis of structural shape and chemical complementarity to GlmU was ranked, based on S value and Lipinski’s drug likeliness in MOE.

Ones with least S-value and Lipinski’s drug likeliness=1 were considered as best inhibitors. Results of GlmU binding with compounds from group I, II and II showed several interactions (Fig. 3, Table 2). 4,4’-[(1E)-5-Hydroxy-4-(methoxymethyl)-1-pentene-1,5-diyl]diphenol from group I, apigenin-7-O-beta-guconopyranoside (also commonly known as apigetrin) from group II and methyl rosmarinate was obtained as the best anti-microbial against *Mycobacterium chelonae* from group III compounds. Common source of 4,4’-[(1E)-5-Hydroxy-4-(methoxymethyl)-1-pentene-1,5-diyl]diphenol is *Alpinia galanga* while apigetrin is sourced from dandelion coffee and *Teucrium gnaphalodes*. Methyl rosmarinate is a constituent of *Rabdosia serra, Ehretia thyrsiflora, Dracocephalum heterophyllum, Perilla frutescens, Nepeta deflersiana, Cordia sinensis* and *Hyptis atrorubens* Poit. ADME and toxicity parameters were also calculates and it was found that apigenin-7-O-beta-gluconopyranoside had lowest blood brain penetration but predicted as a mutagen while methyl rosmarinate and 4,4’-[(1E)-5-Hydroxy-4-(methoxymethyl)-1-pentene-1,5-diyl]diphenol were found to be non-mutagenic and had slightly higher blood brain barrier penetration (Table 3). Analogs of apigenin have shown activity against Mycobacterial species (Tsouh Fokou *et al.*, 2015; Evina *et al.*, 2017) while other compounds have not been reported in literature as anti-mycobacterials yet.

**Fig. 3.**
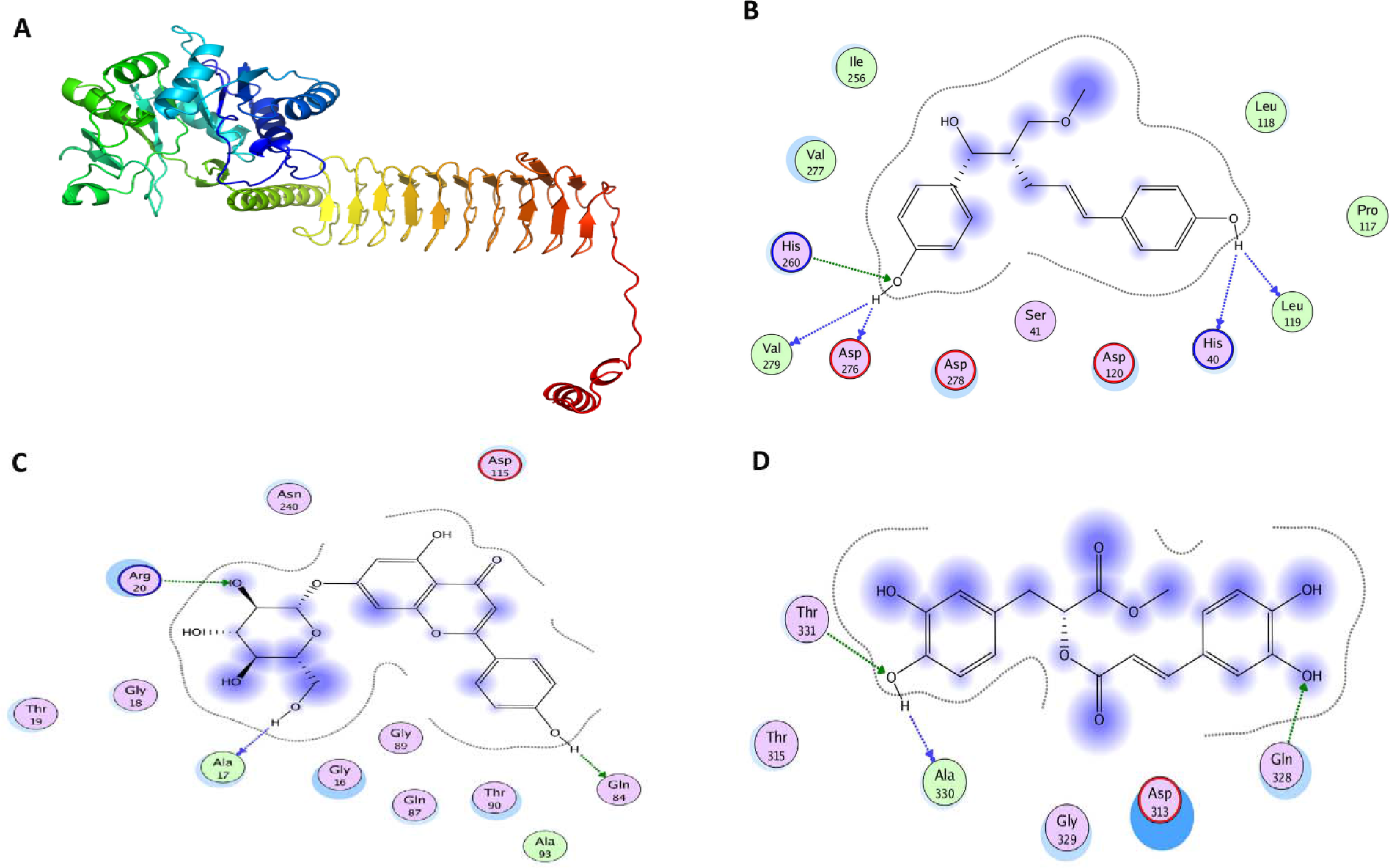
**(A)** Predicted 3D structure of GlmU **(B)** Docked GlmU with 4,4’-[(1E)-5-Hydroxy-4-(methoxymethyl)-1-pentene-1,5-diyl]diphenol showing residue interactions (2D conformation) **(C)** apigenin-7-O-beta-guconopyranoside and GlmU docked complex(2D conformation) **(D)** Methyl rosmarinate and GlmU docked complex(2D conformation). Blue arrowhead represents backbone donor/acceptor while green arrowhead depicts sidechain donor/acceptor, dotted line represents proximity contour.

**Table 2.**
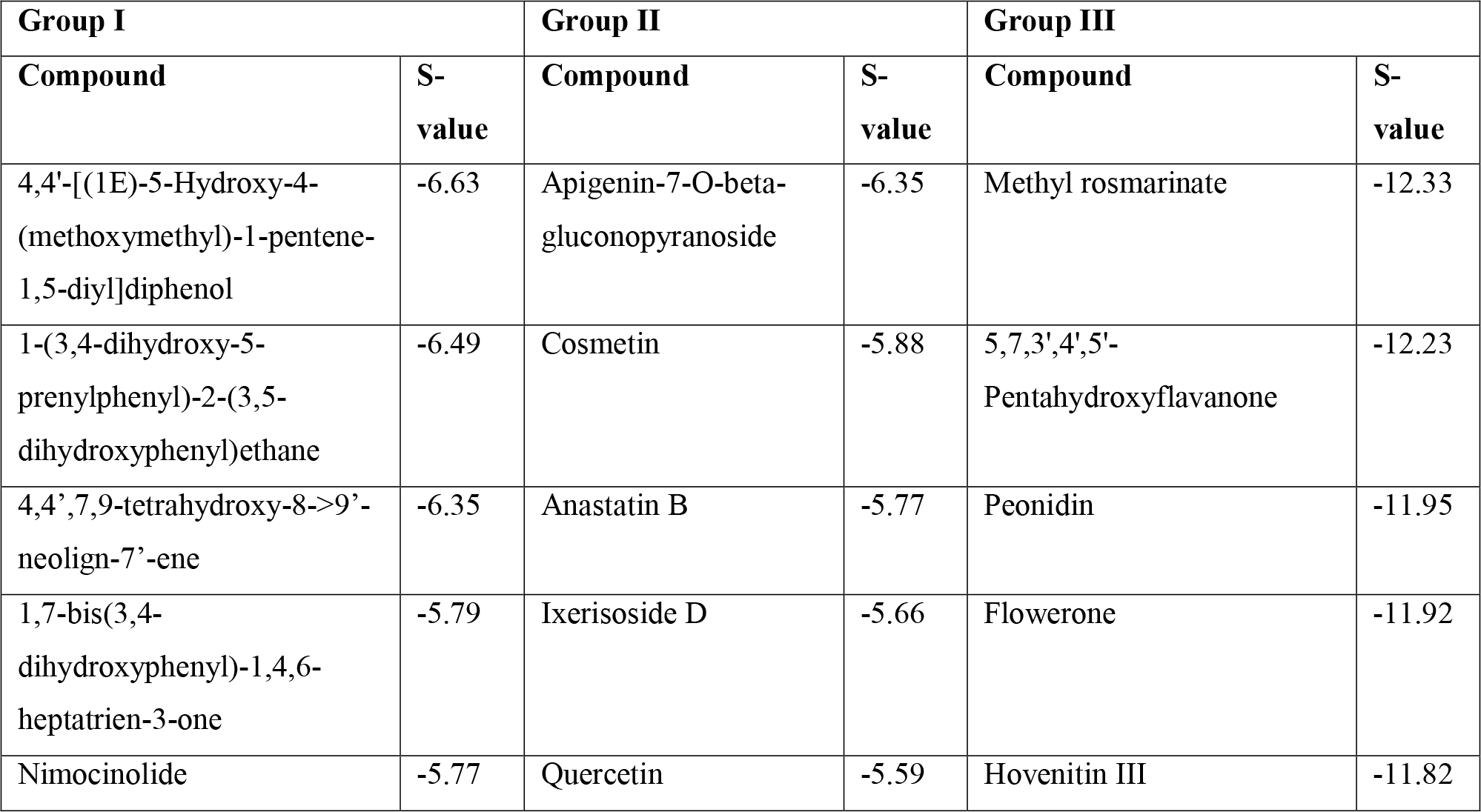
Top 5 compounds against GlmU of *Mycobacterium chelonae* (from each docked group obtained with best inhibition score).

**Table 3.** ADME and toxicity profile of top drug-like molecules from studied phytochemical group compounds.

| Parameter | 4,4'-[(1E)-5-Hydroxy-4-(methoxymethyl)-1-pentene-1,5-diyl]diphenol | Apigenin-7-O-beta-gluconopyranoside | Methyl rosmarinate |
| --- | --- | --- | --- |
| *In vivo* blood-brain barrier penetration | 1.90 | 0.03 | 0.17 |
| CYP450 2D6 inhibition | None | None | None |
| Human intestinal absorption (%) | 90.47 | 47.10 | 76.88 |
| Plasma protein binding (%) | 89.08 | 73.43 | 86.57 |
| Ames test | Non-mutagen | mutagen | Non-mutagen |
| Skin permeability (logKp, cm/hour) | -2.79 | -4.52762 | -3.30 |
| hERG inhibition | Medium risk | High risk | Medium risk |
| Carcinogenicity in mouse | Negative | Positive | Negative |
| Carcinogenicity in rat | Negative | Negative | positive |

We focused on phytochemical screening against *Mycobacterium chelonae* GlmU because plant derived/natural compounds could be used as antibacterial therapeutics for treatment of bacterial infections. Further Lab testing is proposed to know about minimum inhibitory concentration value and other parameters for these compounds against *Mycobacterium chelonae.*

## CONCLUSION

In this study, we proposed GlmU as one of the important therapeutic drug targets for *Mycobacterium chelonae* as it is bifunctional, essential protein for pathogen and has no homology with human proteome. We have provided putative model for phytotherapy against *Mycobacterium chelonae* through virtual screening-based identification of potent metabolites from three data sets, comprising of more than 10,000 compounds. This study could be taken as an initiative for screening and quick designing of phytotherapy against microbes using a computational *modus operandi*. It is expected that our study will also facilitate selection and screening of other *Mycobacterium chelonae* therapeutic target proteins against phytochemical and other relevant compounds for Lab testing and successful entry into drug design pipeline.

## ACKNOWLEDGEMENTS

The authors are thankful to Dr. Waqasuddin Khan (Jamil-ur-Rahman Center for Genome Research, PCMD, ICCBS, University of Karachi) for useful comments and insights that facilitated the analysis. The authors are also thankful to Dr. Alexandra Elbakyan for help with

---

**Supplementary Table 1.** List of bacterial species used for essential proteome prediction of *Mycobacterium chelonae*.

**Supplementary Table 2.** Essential proteome of *Mycobacterium chelonae.*

**Supplementary Table 3.** Non-homologous core proteome of *Mycobacterium chelonae*.

## REFERENCES

Abadio, A. K. R., Kioshima, E. S., Teixeira, M. M., Martins, N. F., Maigret, B. & Felipe, M. S. (2011). Comparative genomics allowed the identification of drug targets against human fungal pathogens. BMC Genomics, 12(1),1.

Aiello, D., Barnes, M. H., Biswas, E. E., Biswas, S. B., Gu, S., Williams, J. D., Bowlin, T. L. & Moir, D. T. (2009). Discovery, characterization and comparison of inhibitors of Bacillus anthracis and Staphylococcus aureus replicative DNA helicases. Bioorganic and Medical Chemistry, 17(13), 4466–76.

Amineni, U., Pradhan, D. & Marisetty, H. (2010) In silico identification of common putative drug targets in Leptospira interrogans. Journal of Chemical Biology, 3, 165–73.

Anthony, K. G., Strych, U., Yeung, K. R., Shoen, C. S., Perez, O., Krause, K. L., Cynamon, M. H., Aristoff, P. A. & Koski, R. A. (2011). New classes of alanine racemase inhibitors identified by high-throughput screening show antimicrobial activity against Mycobacterium tuberculosis. PLoS One, 6(5), e20374.

Azam, S.S. & Shamim, A. (2014). An insight into the exploration of druggable genome of Streptococcus gordonii for the identification of novel therapeutic candidates. Genomics, 104(3), 203–14.

Basharat, Z., Zaib, S. & Yasmin A. (2017). Computational study of some amoebicidal phytochemicals against heat shock protein of Naegleria fowleri. Gene Reports, 1(6), 158–62.

Biasini, M., Bienert, S., Waterhouse, A., Arnold, K., Studer, G., Schmidt, T., Kiefer, F., Cassarino, T.G., Bertoni, M., Bordoli, L. & Schwede, T. (2014). SWISS-MODEL: modelling protein tertiary and quaternary structure using evolutionary information. Nucleic Acids Research, 42(W1), W252–W258.

Bhave, D. P., Wilson III, B., & Carroll, K. S. (2007). Drug targets in mycobacterial sulfur metabolism. Infectious Disorders-Drug Targets, 7(2), 140–158.

Brown-Elliott, B.A., Wallace Jr, R.J., Blinkhorn, R., Crist, C.J. & Mann, L.B. (2001). Successful treatment of disseminated Mycobacterium chelonae infection with linezolid. Clinical Infectious Diseases, 33(8),1433–4.

Butt, A. M., Tahir, S., Nasrullah, I., Idrees, M., Lu, J. & Tong, Y. (2012). Mycoplasma genitalium: A comparative genomics study of metabolic pathways for the identification of drug and vaccine targets. Infection Genetics and Evolution, 12, 53–62.

Choi, Y.J., Oh, E.K., Lee, M.J., Kim, N., Choung, H.K., Khwarg, S.I. (2018). Atypical mycobacterial infection in anophthalmic sockets with porous orbital implant exposure. American Journal of Ophthalmology, In press.

Churgin, D.S., Tran, K.D., Gregori, N.Z., Young, R.C., Alabiad, C. & Flynn Jr H.W. (2018). Multi-drug resistant Mycobacterium chelonae scleral buckle infection. American Journal of Ophthalmology Case Reports, 10:276–8.

Duffield, M., Cooper, I., McAlister, E., Bayliss, M., Ford, D. & Oyston, P. (2010). Predicting conserved essential genes in bacteria: in silico identification of putative drug targets. Molecular Biosystems, 6, 2482–89.

Evina, J.N., Bikobo, D.S., Zintchem, A.A., Nyemeck II, Mbabi, N., Ndedi, E.D., Diboué, P.H., Nyegue, M.A., Atchadé, A.D., Pegnyemb, D.E. & Koert, U. (2017). In vitro antitubercular activity of extract and constituents from the stem bark of Disthemonanthus benthamianus. Revista Brasileira de Farmacognosia, 27(6), 739–43.

Gonzalez-Santiago, T. M. & Drage, L. A. (2015). Nontuberculous Mycobacteria: Skin and soft tissue infections. Dermatologic Clinics, 33(3), 563–577.

Hay, R. J. (2009). Mycobacterium chelonae -a growing problem in soft tissue infection. Current Opinion in Infectious Diseases, 22, 99–101.

Jaén-Luchoro, D., Salvà-Serra, F., Aliaga-Lozano, F., Seguí, C., Busquets, A., Ramírez, A., Ruíz, M., Gomila, M., Lalucat, J. & Bennasar-Figueras, A. (2016). Complete genome sequence of Mycobacterium chelonae type strain CCUG 47445, a rapidly growing species of nontuberculous mycobacteria. Genome Announcements, 4(3), e00550–16.

Jarlier VI, Gutmann, L. & Nikaido, H. (1991). Interplay of cell wall barrier and beta-lactamase activity determines high resistance to beta-lactam antibiotics in Mycobacterium chelonae. Antimicrobial Agents and Chemotherapy. 35(9), 1937–9.

Kosinska, U., Carnrot, C., Eriksson, S., Wang, L. & Eklund, H. (2005). Structure of the substrate complex of thymidine kinase from Ureaplasma urealyticum and investigations of possible drug targets for the enzyme. FEBS Journal, 272(24), 6365–6372.

Lagunin, A.A., Druzhilovsky, D.S., Rudik, A.V., Filimonov, D.A., Gawande, D., Suresh, K., Goel, R., Poroikov, V.V. (2015). Computer evaluation of hidden potential of phytochemicals of medicinal plants of the traditional Indian ayurvedic medicine. Biomeditsinskaia Khimiia, 61(2), 286–97.

Lee, J. Y., Jeong, K. W., Shin, S., Lee, J. U. & Kim, Y. (2012). Discovery of novel selective inhibitors of Staphylococcus aureus-β-ketoacyl acyl carrier protein synthase III. European Journal of Medical Chemistry, 47, 261–269.

Li, Y., Wong, Y. L., Ng, F. M., Liu, B., Wong, Y. X., Poh, Z. Y., Liu, S., Then, S. W., Lee, M. Y., Ng, H. Q. & Huang, Q. (2016). Escherichia coli topoisomerase IV E subunit and an inhibitor binding mode revealed by NMR spectroscopy. Journal of Biological Chemistry, 291(34), 17743–17753.

Luckner, S. R., Liu, N., Am Ende, C.W., Tonge, P.J. & Kisker, C. (2010). A slow, tight binding inhibitor of InhA, the enoyl-acyl carrier protein reductase from Mycobacterium tuberculosis. Journal of Biological Chemistry, 285(19), 14330–14337.

Mehra, R., Rani, C., Mahajan, P., Vishwakarma, R. A., Khan, I. A & Nargotra, A. (2016). Computationally guided identification of novel Mycobacterium tuberculosis GlmU inhibitory leads, their optimization and in vitro validation. ACS Combinatorial Science, 18(2), 100–116.

Mehra, R., Sharma, R., Khan, I. A. & Nargotra, A. (2015). Identification and optimization of Escherichia coli GlmU inhibitors: an in silico approach with validation thereof. European Journal of Medical Chemistry, 92, 78–90.

Mills, S. D., Eakin, A. E., Buurman, E. T., Newman, J. V., Gao, N., Huynh, H., Johnson, K.D., Lahiri, S., Shapiro, A. B., Walkup, G. K. & Yang, W. (2011). Novel bacterial NAD+-dependent DNA ligase inhibitors with broad-spectrum activity and antibacterial efficacy in vivo. Antimicrobial Agents Chemotherapy, 55(3),1088–1096.

Mochalkin, I., Lightle, S., Narasimhan, L., Bornemeier, D., Melnick, M., VanderRoest, S., McDowell L (2008). Structure of a small-molecule inhibitor complexed with GlmU from Haemophilus influenzae reveals an allosteric binding site. Protein Science, 17, 577–82.

Moraes, G. L., Gomes, G. C., deSousa, P. R., Alves, C. N., Govender, T., Kruger, H. G., Maguire, G. E., Lamichhane, G. & Lameira, J. (2015). Structural and functional features of enzymes of Mycobacterium tuberculosis peptidoglycan biosynthesis as targets for drug development. Tuberculosis, 95, 95–111.

Mumtaz, A., Ashfaq, U. A., ulQamar, M. T., Anwar, F., Gulzar, F., Ali, M. A., Saari, N. & Pervez, M. T. (2016). MPD3: a useful medicinal plants database for drug designing. Natural Product Research, 1–9.

Nancy, Y. Y., Wagner, J. R., Laird, M. R., Melli, G., Rey, S., Lo, R., Dao, P., Sahinalp, S. C., Ester, M., Foster, L. J. & Brinkman, F. S. (2010). PSORTb 3.0: improved protein subcellular localization prediction with refined localization subcategories and predictive capabilities for all prokaryotes. Bioinformatics, 26, 1608–15.

Ntie-Kang, F., Telukunta, K.K., Do◻ring, K., Simoben, C.V.A., Moumbock, A.F., Malange, Y.I., Njume, L.E., Yong, J.N., Sippl, W. & Gu◻nther, S. (2017). NANPDB: A resource for natural products from Northern African sources. Journal of Natural Products. 80(7), 2067–76.

Park, B., Awasthi, D., Chowdhury, S. R., Melief, E. H., Kumar, K., Knudson, S. E., Slayden, A. & Ojima, I. (2014). Design, synthesis and evaluation of novel 2, 5, 6-trisubstituted benzimidazoles targeting FtsZ as antitubercular agents. Bioorganic and Medicinal Chemistry, 22(9), 2602–2612.

Patin, D., Bayliss, M., Lecreulx, D. M., Oyston, P. & Blanot, D. (2015). Purification and biochemical characterisation of GlmU from Yersinia pestis. Archives Microbiology, 197(3),371–78.

Patnaik, S., Mohanty, I., Panda, P., Sahu, S. & Dash, M. (2013). Disseminated Mycobacterium chelonae infection: Complicating a case of hidradenitis suppurativa. Indian Dermatology Online Journal, 4(4), 336–396.

Rengarajan, J., Sassetti, C. M., Naroditskaya, V., Sloutsky, A., Bloom, B. R., & Rubin, E. J. (2004). The folate pathway is a target for resistance to the drug para-aminosalicylic acid (PAS) in mycobacteria. Molecular Microbiology, 53(1), 275–282.

Sarangi, A. N., Lohani, M. & Aggarwal, R. (2015). Proteome mining for drug target identification in Listeria monocytogenes EGD-e and structure based virtual screening of a candidate drug target penicillin binding protein. Journal of Microbiological Methods, 111, 9–18.

Shanmugam, A. & Natarajan, J. (2010). Computational genome analyses of metabolic enzymes in Mycobacterium leprae for drug target identification. Bioinformation, 4, 392–95.

Shanmugham, B. & Pan, A. (2013). Identification and characterization of potential therapeutic candidates in emerging human pathogen Mycobacterium abscessus: A novel hierarchical in silico approach. PLOS One, 8(3), 59126.

Sharma, R., Lambu, M. R., Jamwa, U., Rani, C., Chib, R. & Wazir, P. (2016). Escherichia coli N-Acetylglucosamine-1-Phosphate-Uridyltransferase/Glucosamine-1-Phosphate-Acetyltransferase (GlmU) inhibitory activity of terreic acid isolated from Aspergillus terreus. Journal of Biomolecular Screening, 1–12.

Shi, G., Blaszczyk, J., Ji, X. & Yan, H. (2001). Bisubstrate analogue inhibitors of 6-hydroxymethyl-7, 8-dihydropterin pyrophosphokinase: synthesis and biochemical and crystallographic studies. Journal of Medicinal Chemistry, 44(9), 1364–1371.

Slayden, R. A. & Belisle, J. T. (2009). Morphological features and signature gene response elicited by inactivation of FtsI in Mycobacterium tuberculosis. Journal of Antimicrobial Chemotherapy, 63(3), 451–457.

Soares da Costa, T. P., Tieu, W., Yap, M. Y., Zvarec, O., Bell, J. M., Turnidge, J.D., Wallace, J. C., Booker, G. W, Wilce, M. C., Abell, A. D. & Polyak, S.W. (2012). Biotin analogues with antibacterial activity are potent inhibitors of biotin protein ligase. ACS Medicinal Chemistry Letters, 3(6), 509–514.

Strzyz, P. (2016). Protein translocation channelling the route for ER misfolded proteins. Nature Reviews Molecular Cell Biology, 17, 462–463.

Tholander, F. & Sjöberg, B. M. (2012). Discovery of antimicrobial ribonucleotide reductase inhibitors by screening in microwell format. Proceedings of the National Academy of Sciences, 109(25), 9798–9803.

Tortoli, E. (2014). Microbiological features and clinical relevance of new species of the genus Mycobacterium. Clinical Microbiology Reviews, 27(4),727–752.

Tsirigos, K. D., Peters, C., Shu, N., Käll, L. & Elofsson, A. (2015). The TOPCONS web server for consensus prediction of membrane protein topology and signal peptides. Nucleic Acids Research, 485

Tsouh Fokou, P. V., Nyarko, A. K., Appiah-Opong, R., Yamthe, T., Rachel, L., Ofosuhene, M., & Boyom, F. F. (2015). Update on medicinal plants with potency on Mycobacterium ulcerans. BioMed Research International, 917086, 1–16.

Wang, Z., Sun, H., Yao, X., Li, D., Xu, L., Li, Y., Tian, S. & Hou, T. (2016). Comprehensive evaluation of ten docking programs on a diverse set of protein–ligand complexes: the prediction accuracy of sampling power and scoring power. Journal of Physical Chemistry, 18, 12964–75.

Wei, W., Ning, L. W., Ye, Y. N. & Guo, F. B. (2013). Geptop: A Gene Essentiality Prediction Tool for Sequenced Bacterial Genomes Based on Orthology and Phylogeny. PLOS One, 8(8), 72343.

Zhang, Y. & Skolnick, J. (2005). TM-align: a protein structure alignment algorithm based on the TM-score. Nucleic Acids Research, 33(7), 2302–2309.

Zhao, Y., Bacher, A., Illarionov, B., Fischer, M., Georg, G., Ye, Q. Z., Fanwick, P. E., Franzblau, S. G., Wan, B. & Cushman, M. (2009). Discovery and development of the covalent hydrates of trifluoromethylated pyrazoles as riboflavin synthase inhibitors with antibiotic activity against Mycobacterium tuberculosis. Journal of Organic Chemistry, 74(15),5297–5303.

Zhao, Y., Shadrick, W. R., Wallace, M. J., Wu, Y., Griffith, E.C., Qi, J., Yun, M. K., White, W. & Lee, R. E. (2016). Pterin–sulfa conjugates as dihydropteroate synthase inhibitors and antibacterial agents. Bioorganic and Medicinal Chemistry Letters, 26(16), 3950–3954.

Zhuo, H. S., Ni, H. H., Yun, M. Q., He, M. M., Liang, Z. M., Dai Hao Fu, J. Y., Hai, W. Q. & Xing, Z. Y. (2015). The phytochemicals with antagonistic activities toward pathogens of a disease complex caused by Meloidogyne incognita and Ralstonia solanacearum. Journal of Pure and Applied Microbiology, 9, 209–13.

