## Supplementary data for "Drug Target Identification and Virtual Screening in Pursuit of Phytochemical Intervention of *Mycobacterium chelonae*"

Supplementary Table 1. List of bacterial species used for essential proteome prediction of *Mycobacterium chelonae.*

| S. no. | Strain |
| --- | --- |
| 1. | *Acinetobacter baylyi* ADP1 |
| 2. | *Bacillus subtilis* 168 |
| 3. | *Caulobacter crescentus* NA1000 |
| 4. | *Escherichia coli* MG1655 |
| 5. | *Fransicella noicida* U112 |
| 6. | *Haemophilus influenza* Rd KW20 |
| 7. | *Helicobacter pylori* 26695 |
| 8. | *Mycobacterium tuberculosis* H37Rv |
| 9. | *Mycoplasma genitalium* G37 |
| 10. | *Mycoplasma pulmonis* UAB CTIP |
| 11. | *Pseudomonas aeruginosa* UCBPP-PA14 |
| 12. | *Salmonella typhi* Ty2 |
| 13. | *Salmonella typhimurium* LT2 |
| 14. | *Staphylococcus aureus* N315 |
| 15. | *Staphylococcus aureus* NCTC 8325 |
| 16. | *Streptococcus pneumonia* R6 |
| 17. | *Streptococcus pneumonia* TIGR4 |
| 18. | *Streptococcus sangunis* SK36 |
| 19. | *Vibrio cholera* N166961 |

Supplementary Table 2. Essential proteome of *Mycobacterium chelonae.*

| S. no. | Essentiality Score | Protein ID |
| --- | --- | --- |
| 1. | 0.3967 | WP_003418601.1 |
| 2. | 0.3967 | WP_003883485.1 |
| 3. | 0.4132 | WP_003929602.1 |
| 4. | 0.4132 | WP_005055645.1 |
| 5. | 0.4133 | WP_005055680.1 |
| 6. | 0.3622 | WP_005055719.1 |
| 7. | 0.3968 | WP_005056050.1 |
| 8. | 0.1802 | WP_005056735.1 |
| 9. | 0.3599 | WP_005058700.1 |
| 10. | 0.304 | WP_005060346.1 |
| 11. | 0.2645 | WP_005063156.1 |
| 12. | 0.4132 | WP_005077598.1 |
| 13. | 0.3777 | WP_030093446.1 |
| 14. | 0.2024 | WP_030093470.1 |
| 15. | 0.3438 | WP_030093498.1 |
| 16. | 0.2245 | WP_030093515.1 |
| 17. | 0.1799 | WP_030093516.1 |
| 18. | 0.2457 | WP_030093559.1 |
| 19. | 0.1568 | WP_030093563.1 |
| 20. | 0.3063 | WP_030093572.1 |
| 21. | 0.2433 | WP_030093606.1 |
| 22. | 0.1774 | WP_030093611.1 |
| 23. | 0.2869 | WP_030093878.1 |
| 24. | 0.3776 | WP_030093885.1 |
| 25. | 0.3041 | WP_030093922.1 |
| 26. | 0.3795 | WP_030093923.1 |
| 27. | 0.4131 | WP_030093948.1 |
| 28. | 0.445 | WP_030094002.1 |
| 29. | 0.4293 | WP_030094003.1 |
| 30. | 0.3966 | WP_030094010.1 |
| 31. | 0.3621 | WP_030094034.1 |
| 32. | 0.3599 | WP_030094038.1 |
| 33. | 0.1771 | WP_030094131.1 |
| 34. | 0.2435 | WP_030094135.1 |
| 35. | 0.2028 | WP_030094777.1 |
| 36. | 0.2028 | WP_030094835.1 |
| 37. | 0.3255 | WP_030094894.1 |
| 38. | 0.3063 | WP_030094903.1 |
| 39. | 0.344 | WP_030094907.1 |
| 40. | 0.3796 | WP_030095128.1 |
| 41. | 0.344 | WP_030095153.1 |
| 42. | 0.3966 | WP_030095371.1 |
| 43. | 0.1802 | WP_030095414.1 |
| 44. | 0.1803 | WP_030095429.1 |
| 45. | 0.2436 | WP_030095452.1 |
| 46. | 0.1542 | WP_030095583.1 |
| 47. | 0.3439 | WP_030095704.1 |
| 48. | 0.3255 | WP_030095787.1 |
| 49. | 0.1774 | WP_030095815.1 |
| 50. | 0.2867 | WP_030095824.1 |
| 51. | 0.2434 | WP_030095840.1 |
| 52. | 0.1801 | WP_030095982.1 |
| 53. | 0.2644 | WP_030095993.1 |
| 54. | 0.1569 | WP_030096236.1 |
| 55. | 0.2667 | WP_030096504.1 |
| 56. | 0.3065 | WP_030096568.1 |
| 57. | 0.3966 | WP_030096617.1 |
| 58. | 0.3418 | WP_030096618.1 |
| 59. | 0.4132 | WP_030096639.1 |
| 60. | 0.4451 | WP_030096640.1 |
| 61. | 0.4294 | WP_030096665.1 |
| 62. | 0.445 | WP_030096693.1 |
| 63. | 0.445 | WP_030096694.1 |
| 64. | 0.4293 | WP_030096695.1 |
| 65. | 0.4133 | WP_030096696.1 |
| 66. | 0.3966 | WP_030096697.1 |
| 67. | 0.4294 | WP_030096699.1 |
| 68. | 0.4133 | WP_030096705.1 |
| 69. | 0.4132 | WP_030096712.1 |
| 70. | 0.4451 | WP_030096713.1 |
| 71. | 0.4294 | WP_030096714.1 |
| 72. | 0.4293 | WP_030096715.1 |
| 73. | 0.4114 | WP_030096716.1 |
| 74. | 0.4293 | WP_030096726.1 |
| 75. | 0.4113 | WP_030096727.1 |
| 76. | 0.3947 | WP_030096736.1 |
| 77. | 0.4451 | WP_030096737.1 |
| 78. | 0.4294 | WP_030096756.1 |
| 79. | 0.2459 | WP_030096781.1 |
| 80. | 0.344 | WP_030096989.1 |
| 81. | 0.3439 | WP_030097167.1 |
| 82. | 0.4293 | WP_030097637.1 |
| 83. | 0.3418 | WP_030097640.1 |
| 84. | 0.3794 | WP_030097693.1 |
| 85. | 0.4132 | WP_030097698.1 |
| 86. | 0.1773 | WP_030097761.1 |
| 87. | 0.4451 | WP_030098010.1 |
| 88. | 0.3599 | WP_030098059.1 |
| 89. | 0.2667 | WP_030098083.1 |
| 90. | 0.1771 | WP_043076460.1 |
| 91. | 0.2665 | WP_044103925.1 |
| 92. | 0.1541 | WP_044104001.1 |
| 93. | 0.1569 | WP_044104190.1 |
| 94. | 0.18 | WP_044104593.1 |
| 95. | 0.3797 | WP_044104607.1 |
| 96. | 0.2433 | WP_044105188.1 |
| 97. | 0.4603 | WP_044105291.1 |
| 98. | 0.3599 | WP_046251964.1 |
| 99. | 0.445 | WP_046251970.1 |
| 100. | 0.2218 | WP_046252008.1 |
| 101. | 0.4603 | WP_046252123.1 |
| 102. | 0.1772 | WP_046252174.1 |
| 103. | 0.246 | WP_046252207.1 |
| 104. | 0.2221 | WP_046252249.1 |
| 105. | 0.1568 | WP_046252257.1 |
| 106. | 0.3439 | WP_046252353.1 |
| 107. | 0.3439 | WP_046252392.1 |
| 108. | 0.2867 | WP_046252393.1 |
| 109. | 0.2869 | WP_046252394.1 |
| 110. | 0.222 | WP_046252395.1 |
| 111. | 0.2027 | WP_046252399.1 |
| 112. | 0.3796 | WP_046252400.1 |
| 113. | 0.2868 | WP_046252415.1 |
| 114. | 0.4293 | WP_046252416.1 |
| 115. | 0.1541 | WP_046252467.1 |
| 116. | 0.1541 | WP_046252468.1 |
| 117. | 0.2435 | WP_046252502.1 |
| 118. | 0.1569 | WP_046252531.1 |
| 119. | 0.1542 | WP_046252630.1 |
| 120. | 0.3061 | WP_046252725.1 |
| 121. | 0.1799 | WP_046252740.1 |
| 122. | 0.3795 | WP_046252754.1 |
| 123. | 0.3256 | WP_046252760.1 |
| 124. | 0.2845 | WP_046252764.1 |
| 125. | 0.4293 | WP_046252765.1 |
| 126. | 0.1541 | WP_046252766.1 |
| 127. | 0.3795 | WP_046252768.1 |
| 128. | 0.3441 | WP_046252769.1 |
| 129. | 0.3439 | WP_046252782.1 |
| 130. | 0.1538 | WP_046252806.1 |
| 131. | 0.2219 | WP_046252830.1 |
| 132. | 0.3233 | WP_046252838.1 |
| 133. | 0.2643 | WP_046252951.1 |
| 134. | 0.2666 | WP_046252969.1 |
| 135. | 0.1799 | WP_046252971.1 |
| 136. | 0.4293 | WP_046252972.1 |
| 137. | 0.2219 | WP_046252974.1 |
| 138. | 0.2459 | WP_046252975.1 |
| 139. | 0.3064 | WP_046252976.1 |
| 140. | 0.2867 | WP_046252977.1 |
| 141. | 0.2026 | WP_046252978.1 |
| 142. | 0.36 | WP_046252999.1 |
| 143. | 0.2844 | WP_046253019.1 |
| 144. | 0.4293 | WP_046253176.1 |
| 145. | 0.3064 | WP_046253177.1 |
| 146. | 0.1544 | WP_046253179.1 |
| 147. | 0.3967 | WP_046253183.1 |
| 148. | 0.3966 | WP_046253184.1 |
| 149. | 0.3253 | WP_046253193.1 |
| 150. | 0.2024 | WP_046253230.1 |
| 151. | 0.3966 | WP_046253234.1 |
| 152. | 0.1565 | WP_046253258.1 |
| 153. | 0.3794 | WP_046253264.1 |
| 154. | 0.2664 | WP_046253323.1 |
| 155. | 0.2246 | WP_046253374.1 |
| 156. | 0.1569 | WP_046253375.1 |
| 157. | 0.2433 | WP_046253395.1 |
| 158. | 0.2456 | WP_046253411.1 |
| 159. | 0.362 | WP_046253413.1 |
| 160. | 0.3777 | WP_046253414.1 |
| 161. | 0.4134 | WP_046253415.1 |
| 162. | 0.4133 | WP_046253416.1 |
| 163. | 0.3796 | WP_046253417.1 |
| 164. | 0.287 | WP_046253418.1 |
| 165. | 0.4295 | WP_046253419.1 |
| 166. | 0.2246 | WP_046253420.1 |
| 167. | 0.445 | WP_046253421.1 |
| 168. | 0.2003 | WP_046253513.1 |
| 169. | 0.3621 | WP_046253587.1 |
| 170. | 0.4293 | WP_046253608.1 |
| 171. | 0.1775 | WP_046253619.1 |
| 172. | 0.4131 | WP_046253621.1 |
| 173. | 0.3966 | WP_046253625.1 |
| 174. | 0.3795 | WP_046253628.1 |
| 175. | 0.3795 | WP_046253633.1 |
| 176. | 0.2243 | WP_046253734.1 |
| 177. | 0.1797 | WP_046253762.1 |
| 178. | 0.344 | WP_046253765.1 |
| 179. | 0.445 | WP_046253813.1 |
| 180. | 0.2667 | WP_046253816.1 |
| 181. | 0.4432 | WP_046253819.1 |
| 182. | 0.2669 | WP_046253834.1 |
| 183. | 0.2644 | WP_046253835.1 |
| 184. | 0.2244 | WP_046253847.1 |
| 185. | 0.2666 | WP_046253856.1 |
| 186. | 0.3252 | WP_046253865.1 |
| 187. | 0.3618 | WP_046253866.1 |
| 188. | 0.2001 | WP_046253878.1 |
| 189. | 0.2667 | WP_046253888.1 |
| 190. | 0.2001 | WP_046253889.1 |
| 191. | 0.4293 | WP_046253891.1 |
| 192. | 0.2221 | WP_046253892.1 |
| 193. | 0.445 | WP_046253897.1 |
| 194. | 0.3797 | WP_046253898.1 |
| 195. | 0.3795 | WP_046253899.1 |
| 196. | 0.2667 | WP_046253910.1 |
| 197. | 0.2026 | WP_046253913.1 |
| 198. | 0.3964 | WP_046253920.1 |
| 199. | 0.2 | WP_046253928.1 |
| 200. | 0.3947 | WP_046253937.1 |
| 201. | 0.3966 | WP_046253941.1 |
| 202. | 0.2435 | WP_046253948.1 |
| 203. | 0.154 | WP_046253949.1 |
| 204. | 0.3965 | WP_046253960.1 |
| 205. | 0.3622 | WP_046254040.1 |
| 206. | 0.2246 | WP_046254047.1 |
| 207. | 0.4133 | WP_046254049.1 |
| 208. | 0.1801 | WP_046254076.1 |
| 209. | 0.1801 | WP_046254077.1 |
| 210. | 0.2459 | WP_046254104.1 |
| 211. | 0.177 | WP_046254111.1 |
| 212. | 0.3775 | WP_046254113.1 |
| 213. | 0.1543 | WP_046254124.1 |
| 214. | 0.4132 | WP_046254138.1 |
| 215. | 0.1775 | WP_046254142.1 |
| 216. | 0.4133 | WP_046254145.1 |
| 217. | 0.2221 | WP_046254166.1 |
| 218. | 0.3234 | WP_046254169.1 |
| 219. | 0.1568 | WP_046254170.1 |
| 220. | 0.3234 | WP_046254171.1 |
| 221. | 0.4294 | WP_046254180.1 |
| 222. | 0.4293 | WP_046254186.1 |
| 223. | 0.2459 | WP_046254195.1 |
| 224. | 0.3795 | WP_046254197.1 |
| 225. | 0.3442 | WP_046254198.1 |
| 226. | 0.4131 | WP_046254206.1 |
| 227. | 0.1569 | WP_046254218.1 |
| 228. | 0.342 | WP_046254220.1 |
| 229. | 0.2461 | WP_046254239.1 |
| 230. | 0.3255 | WP_046254241.1 |
| 231. | 0.445 | WP_046254249.1 |
| 232. | 0.3254 | WP_046254270.1 |
| 233. | 0.2867 | WP_046254272.1 |
| 234. | 0.4603 | WP_046254273.1 |
| 235. | 0.3233 | WP_046254285.1 |
| 236. | 0.3065 | WP_046254286.1 |
| 237. | 0.1539 | WP_046254346.1 |
| 238. | 0.3231 | WP_046254350.1 |
| 239. | 0.1568 | WP_046254399.1 |
| 240. | 0.1773 | WP_046254400.1 |
| 241. | 0.1541 | WP_046254460.1 |
| 242. | 0.4132 | WP_046254491.1 |
| 243. | 0.3233 | WP_046254500.1 |
| 244. | 0.342 | WP_046254518.1 |
| 245. | 0.2667 | WP_046254533.1 |
| 246. | 0.1776 | WP_046254564.1 |
| 247. | 0.1544 | WP_046254565.1 |
| 248. | 0.3965 | WP_046254569.1 |
| 249. | 0.344 | WP_046254576.1 |
| 250. | 0.2026 | WP_046254593.1 |
| 251. | 0.4113 | WP_046254609.1 |
| 252. | 0.3948 | WP_046254611.1 |
| 253. | 0.3439 | WP_046254612.1 |
| 254. | 0.2024 | WP_046254614.1 |
| 255. | 0.3441 | WP_046254617.1 |
| 256. | 0.3256 | WP_046254623.1 |
| 257. | 0.36 | WP_046254636.1 |
| 258. | 0.4293 | WP_046254637.1 |
| 259. | 0.4132 | WP_046254638.1 |
| 260. | 0.4451 | WP_046254654.1 |
| 261. | 0.3967 | WP_046254658.1 |
| 262. | 0.4294 | WP_046254659.1 |
| 263. | 0.2455 | WP_046254679.1 |
| 264. | 0.4131 | WP_046254680.1 |
| 265. | 0.4293 | WP_046254693.1 |
| 266. | 0.3232 | WP_046254709.1 |
| 267. | 0.2869 | WP_046254710.1 |
| 268. | 0.2247 | WP_046254782.1 |
| 269. | 0.3947 | WP_046254838.1 |
| 270. | 0.3063 | WP_046254922.1 |
| 271. | 0.1538 | WP_046254956.1 |
| 272. | 0.3621 | WP_046254994.1 |
| 273. | 0.36 | WP_046255356.1 |
| 274. | 0.3966 | WP_046255370.1 |
| 275. | 0.2868 | WP_046255378.1 |
| 276. | 0.2665 | WP_046255383.1 |
| 277. | 0.1999 | WP_046255384.1 |
| 278. | 0.1568 | WP_046255385.1 |
| 279. | 0.18 | WP_046255391.1 |
| 280. | 0.4114 | WP_046255400.1 |
| 281. | 0.4293 | WP_046255441.1 |
| 282. | 0.2844 | WP_046255563.1 |
| 283. | 0.3966 | WP_046255608.1 |
| 284. | 0.1567 | WP_046255609.1 |
| 285. | 0.18 | WP_046255715.1 |
| 286. | 0.4133 | WP_046255722.1 |
| 287. | 0.4131 | WP_046255749.1 |
| 288. | 0.4132 | WP_046255752.1 |
| 289. | 0.2645 | WP_046255846.1 |
| 290. | 0.3044 | WP_046255877.1 |
| 291. | 0.4275 | WP_046255899.1 |
| 292. | 0.3064 | WP_046255923.1 |
| 293. | 0.2668 | WP_046256002.1 |
| 294. | 0.1801 | WP_046256013.1 |
| 295. | 0.2245 | WP_046256037.1 |
| 296. | 0.1568 | WP_046256048.1 |
| 297. | 0.3775 | WP_046256055.1 |
| 298. | 0.2222 | WP_046256154.1 |
| 299. | 0.3967 | WP_052740089.1 |
| 300. | 0.3968 | WP_052740247.1 |
| 301. | 0.2846 | WP_052740383.1 |
| 302. | 0.3441 | WP_064393527.1 |
| 303. | 0.2242 | WP_064393603.1 |
| 304. | 0.3419 | WP_064393605.1 |
| 305. | 0.2001 | WP_064393607.1 |

Supplementary Table 3. Non-homologous core proteins of *M. chelonae*.

| S. no. | Accession ID |
| --- | --- |
| 1. | WP_005055645 |
| 2. | WP_005055719 |
| 3. | WP_005077598 |
| 4. | WP_030093515 |
| 5. | WP_030093516 |
| 6. | WP_030093559 |
| 7. | WP_030093563 |
| 8. | WP_030093611 |
| 9. | WP_030093923 |
| 10. | WP_030093948 |
| 11. | WP_030094003 |
| 12. | WP_030094034 |
| 13. | WP_030094038 |
| 14. | WP_030094135 |
| 15. | WP_030094907 |
| 16. | WP_030095153 |
| 17. | WP_030095371 |
| 18. | WP_030095414 |
| 19. | WP_030095583 |
| 20. | WP_030095787 |
| 21. | WP_030095815 |
| 22. | WP_030095840 |
| 23. | WP_030095982 |
| 24. | WP_030096618 |
| 25. | WP_030096640 |
| 26. | WP_030096696 |
| 27. | WP_030096697 |
| 28. | WP_030096705 |
| 29. | WP_030096712 |
| 30. | WP_030096714 |
| 31. | WP_030096716 |
| 32. | WP_030096726 |
| 33. | WP_030097637 |
| 34. | WP_030097640 |
| 35. | WP_030097693 |
| 36. | WP_030097698 |
| 37. | WP_030098059 |
| 38. | WP_044104190 |
| 39. | WP_046251964 |
| 40. | WP_046251970 |
| 41. | WP_046252008 |
| 42. | WP_046252249 |
| 43. | WP_046252257 |
| 44. | WP_046252393 |
| 45. | WP_046252394 |
| 46. | WP_046252395 |
| 47. | WP_046252399 |
| 48. | WP_046252415 |
| 49. | WP_046252468 |
| 50. | WP_046252760 |
| 51. | WP_046252764 |
| 52. | WP_046252766 |
| 53. | WP_046252769 |
| 54. | WP_046252806 |
| 55. | WP_046252838 |
| 56. | WP_046252971 |
| 57. | WP_046252974 |
| 58. | WP_046252975 |
| 59. | WP_046252978 |
| 60. | WP_046252999 |
| 61. | WP_046253019 |
| 62. | WP_046253183 |
| 63. | WP_046253264 |
| 64. | WP_046253413 |
| 65. | WP_046253414 |
| 66. | WP_046253415 |
| 67. | WP_046253416 |
| 68. | WP_046253417 |
| 69. | WP_046253418 |
| 70. | WP_046253419 |
| 71. | WP_046253421 |
| 72. | WP_046253513 |
| 73. | WP_046253619 |
| 74. | WP_046253813 |
| 75. | WP_046253816 |
| 76. | WP_046253835 |
| 77. | WP_046253878 |
| 78. | WP_046253888 |
| 79. | WP_046253892 |
| 80. | WP_046253910 |
| 81. | WP_046253928 |
| 82. | WP_046254049 |
| 83. | WP_046254104 |
| 84. | WP_046254169 |
| 85. | WP_046254170 |
| 86. | WP_046254171 |
| 87. | WP_046254186 |
| 88. | WP_046254197 |
| 89. | WP_046254198 |
| 90. | WP_046254241 |
| 91. | WP_046254273 |
| 92. | WP_046254346 |
| 93. | WP_046254350 |
| 94. | WP_046254399 |
| 95. | WP_046254500 |
| 96. | WP_046254518 |
| 97. | WP_046254611 |
| 98. | WP_046254614 |
| 99. | WP_046254636 |
| 100. | WP_046254637 |
| 101. | WP_046254654 |
| 102. | WP_046254658 |
| 103. | WP_046254659 |
| 104. | WP_046254709 |
| 105. | WP_046254710 |
| 106. | WP_046254782 |
| 107. | WP_046254838 |
| 108. | WP_046254956 |
| 109. | WP_046255356 |
| 110. | WP_046255385 |
| 111. | WP_046255722 |
| 112. | WP_046255749 |
| 113. | WP_046255923 |
| 114. | WP_046256048 |
| 115. | WP_046256055 |
| 116. | WP_046256154 |
| 117. | WP_052740089 |
